# Imaging genomics reveals genetic architecture of the globular human braincase

**DOI:** 10.1101/2024.03.20.585712

**Authors:** Barbara Molz, Else Eising, Gökberk Alagöz, Dick Schijven, Clyde Francks, Philipp Gunz, Simon E. Fisher

**Affiliations:** Language and Genetics Department, Max Planck Institute for Psycholinguistics; Nijmegen, The Netherlands; Donders Institute for Brain, Cognition and Behaviour, Radboud University; Nijmegen, The Netherlands; Department of Cognitive Neuroscience, Radboud University Medical Center; Nijmegen, The Netherlands; Department of Human Origins, Max-Planck-Institute for Evolutionary Anthropology; Leipzig, Germany

## Abstract

Compared with our fossil ancestors and Neandertal kin, modern humans have evolved a distinctive skull shape, with a rounder braincase and more delicate face. Competing explanations for this rounder skull have either linked it to changes in brain organisation, or seen it as a by-product of gracilization (evolution of thinner and lighter skeletal anatomy). Here, we combined palaeoanthropological data from hominin fossils and imaging genomics data from living humans to gain insight into evolutionary and developmental mechanisms shaping this uniquely modern human phenotype. We analysed endocranial globularity from magnetic resonance imaging (MRI) brain scans and genetic data of more than 33,000 adults. We discovered 28 genomic loci significantly associated with endocranial globularity. There was genetic overlap with the brain’s ventricular system, white matter microstructure, and sulcal morphology, and with multivariate genetic analyses of reading/language skills, but not with general cognition. The associated genes exhibited enriched expression in the brain during prenatal development and early childhood. The connection to the ventricular system hints at a role for cerebrospinal fluid pressure in shaping the endocranium during development. Genes linked to endocranial globularity also showed enhanced expression in the cardiovascular and female reproductive systems. This finding suggests co-evolutionary pathways whereby changes impacting factors such as energy needs, pregnancy, or fertility concurrently shape the brain and its structure.

---

During foetal and infant development, the growing brain leaves an impression inside the skull. Such endocranial imprints (endocasts) in fossil skulls document evolutionary changes in brain volume, folding patterns, and brain shape. Modern humans have a distinct endocranial shape, that is more globular than the elongated endocrania seen in all other fossil hominins and apes^1,2^. This endocranial globularity involves increased parietal bulging and an enlarged posterior cranial fossa, which houses the cerebellum (Fig. 1A). This unique phenotype emerged within the *Homo sapiens* lineage^3^, and is the most recently evolved anatomical trait of our species^2–5^.

**Fig. 1.**
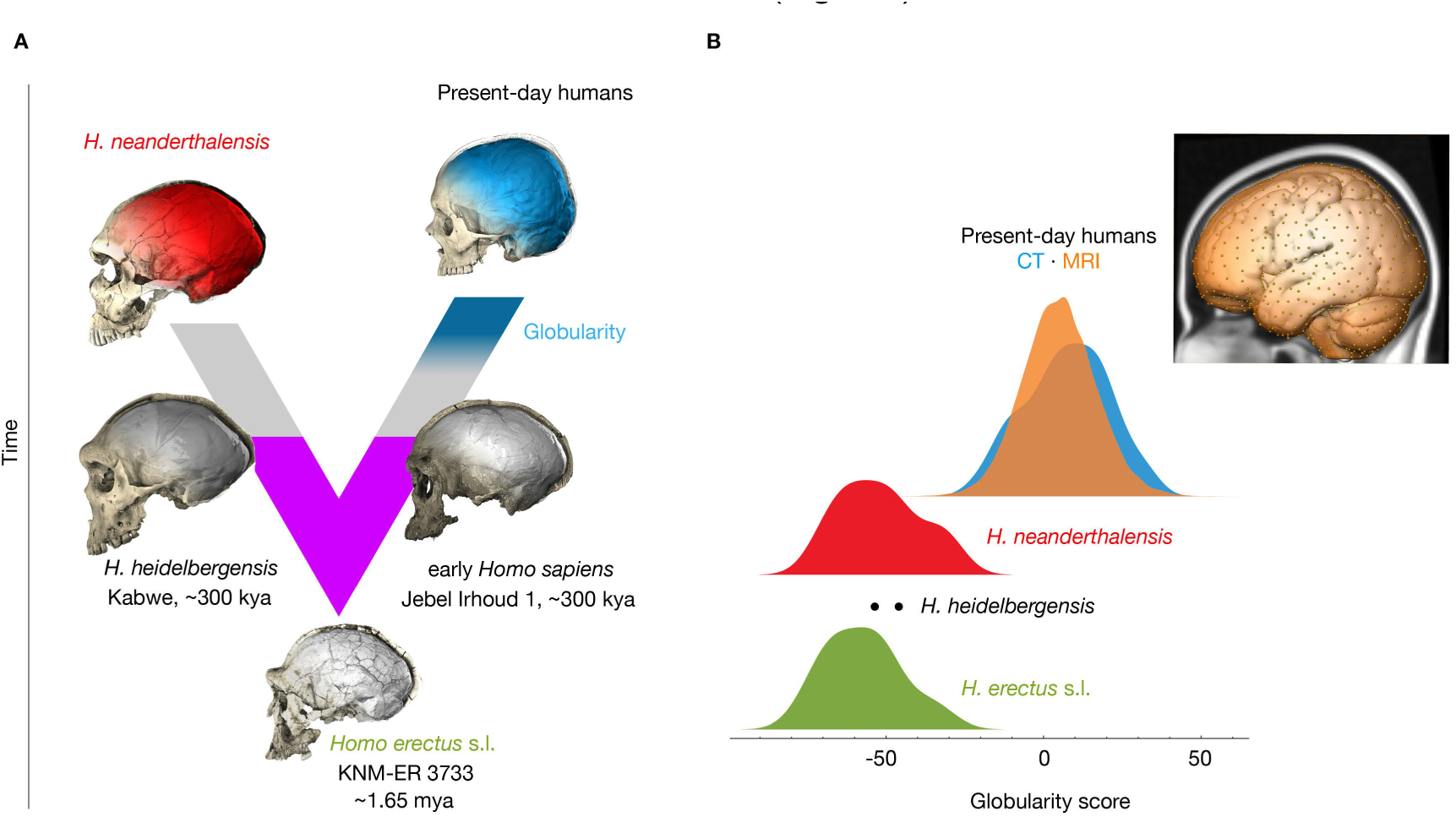
An evolutionarily derived globularity score captures interindividual variability in present-day human endocranial shape. (A) Neandertals and present-day humans share a common ancestor that lived more than 500 kya^12,13^ and likely evolved from *Homo erectus* s.l. in Africa. Purple: Fossils document an increase of endocranial volumes in the lineages leading to Neandertals, and to *Homo sapiens*, respectively. Early *Homo sapiens* like, e.g., Jebel Irhoud 1, have endocranial volumes as large as present-day humans, but elongated endocrania. The emergence of the globularity phenotype in the *Homo sapiens* lineage is symbolized in blue. Virtual endocasts based on CT data are shown for a present-day human, a Neandertal (La Ferrassie 1), *Homo heidelbergensis* (Kabwe), early *Homo sapiens* (Jebel Irhoud 1), and *Homo erectus* (KNM-ER 3733). (B) Histograms of globularity scores computed from different modalities (orange: MRI-based scores of present-day humans, N = 34,595; blue: CT-based scores of present-day humans, N = 89; red: CT-based scores of Neandertals, N = 10; grey: CT-based scores of *H. heidelbergensis*, N = 2; green: CT-based scores of *H. erectus*, N = 8; The endocranial measurement coordinates are shown on the endocast segmentation of the MNI 152 brain template).

Endocranial shape changes are not causally related to evolutionary changes in endocranial volume, since the endocranial volumes of Neandertals and early *Homo sapiens* (∼300 kya)^2,3^ fall within the range of modern variation, but these fossils still have elongated endocrania. In present-day humans, globularity emerges during early ontogeny^6^ – an important period for establishing the white-matter networks of the brain. Consequently, endocranial globularity has been associated with the timing and pattern of brain growth and development, especially the process of myelination^2,5^. Another theory suggests that evolutionary changes in facial size and robusticity, possibly related to diet, were the main drivers of endocranial shape changes within the *Homo sapiens* lineage^7^. However, it is not possible to uncover the underlying developmental processes based on endocasts alone, as these represent only the brain’s outer surface. Here, we combine palaeoanthropology with archaic and neuroimaging genomics, and use interindividual variation in present-day humans as a window into our evolutionary past. We investigate the biological mechanisms underlying endocranial globularity to enhance our understanding of human brain evolution.

An earlier focused study, which used MRI brain scans and DNA data of around 4,500 healthy adults to look specifically for effects of different introgressed Neandertal genetic fragments on endocranial globularity, found associations with the expression of two genes involved in neurogenesis and myelination, respectively^5^. Here, we performed a systematic investigation of the biological bases of endocranial globularity, based on morphometric data from hominin fossil endocasts, integrated with phenotypic and genomic data in a much larger sample.

We used a morphometric analysis of MRI data from lifespan and cross-sectional biobanking cohorts to create a metric that quantifies endocranial globularity, and investigated possible differences across age and sex. We then performed a genome-wide search to identify DNA variants associated with interindividual variation in endocranial globularity in tens of thousands of present-day individuals. Derived summary statistics were used to gain insight into the shared genetic architecture of endocranial globularity and a range of anatomical brain imaging traits. We moved beyond the brain to investigate genetically-mediated associations with disease phenotypes and other complex traits. Finally, we studied the underlying biological mechanisms revealed by enrichment and functional analyses of associated loci, and contributions of genes and genomic regions that show evolutionary changes along the human lineage, on timescales relevant for the emergence of this distinctive human trait.

## An evolutionarily derived quantitative index of present-day human endocranial shape

We used geometric morphometrics^8^ to derive a summary-measure of endocranial globularity from T1-weighted brain images of two independent epidemiological datasets comprising in total 34,595 individuals with European ancestry (Cambridge Centre for Ageing Neuroscience (CamCAN)^9,10^ and UK Biobank (UKB)^11^). We first digitized a mesh of 935 coordinates on virtual endocasts from computed tomographic (CT) scans of fossil and present-day human crania, as well as the MNI 152 brain template. We used Procrustes superimposition to standardize for position, orientation, and scale, and then projected each individual onto the vector between the average shape of Neandertals and present-day humans. This yields a *globularity score*, an evolutionarily derived summary measure of overall endocranial shape^5^. While globularity scores of present-day humans derived from CT scans and structural MRI both show a high degree of interindividual variability, scores from these two modalities cluster together, displaying a minimal overlap with globularity scores of extinct humans, based on fossil data (Fig. 1B).

We first tested for potential effects of ageing on globularity scores using a lifespan dataset of healthy adult participants aged 18-88 years (CamCAN, N = 644), and found no association between age and endocranial globularity (Age: t = 0.536, p = 0.592; Age^2^: t = -0.487, p = 0.626) as well as no interaction of age and sex (Age*Sex: t = -0546, p = 0.585; Age^2^*Sex: t = 0.415, p = 0.678) (Supplementary Fig. 1A). Analysis of globularity scores derived from the UK Biobank (N = 33,951), a large-scale cohort with a limited age-range clustering in the second half of life (mean + SD age 63.77 years ± 7.52), showed a small effect that was statistically significant (Age: t = - 3.822, p = 1.32 x 10^-4^; Age^2^: t =3.709, p =2.08 x 10^-4^) with no significant interaction terms (Age*Sex: t =1.695, p = 0.0.0900; Age^2^*Sex: t = -1.377, p = 0.169), while overall only accounting for 0.4% of the total variance (F(5, 33945) = 25.1, p < 2.2 x 10^-16^, R^2^_Adjusted_ = 0.0035).

We also assessed possible sex effects on endocranial globularity and found a minor but significant difference in globularity scores between males and females in the UK Biobank (t =8.245, p = < 2 x 10^-16^, Cohen’s d = -0.0004) (Supplementary Fig. 1B). Next, we investigated the relationship with estimated total intracranial volume (eTIV) in UK Biobank (N = 33,577), finding a statistically significant but subtle association, where eTIV explained 3.69% of the variance of present-day endocranial globularity scores (F(3, 33573) = 430.1, p < 2.2 x 10^-16^, R^2^_Adjusted_ = 0.0369) (Supplementary Fig. 1C). Lastly, using participant-specific health information available in UK Biobank, we showed that globularity scores derived from people with distinct neuroanatomical or neurodegenerative brain disorders (N = 2,338) do not differ from those of individuals without such a diagnosis (t = -0.039, p = 0.97, Cohen’s d = -0.00084) (Supplementary Fig. 1D).

## Genetics of interindividual differences in endocranial globularity

Next, we performed a genome-wide association study (GWAS) to identify genetic variants associated with interindividual differences in endocranial globularity. As age, neuroanatomical and neurodegenerative disorders, and intracranial brain volume had either no or only marginal influence on globularity scores, all 33,951 UK Biobank participants that passed genetic and phenotypic quality control were included. These analyses identified 28 genome-wide significant loci (p ≤ 5 ×10^−8^), encompassing a total of 155 independent (r^2^ ≤ 0.6) genome-wide significant single-nucleotide polymorphisms (SNPs) and insertions or deletions (indels), associated with interindividual differences in endocranial globularity (Fig. 2A, Supplementary Fig. 2, Supplementary Table 1).

**Fig. 2.**
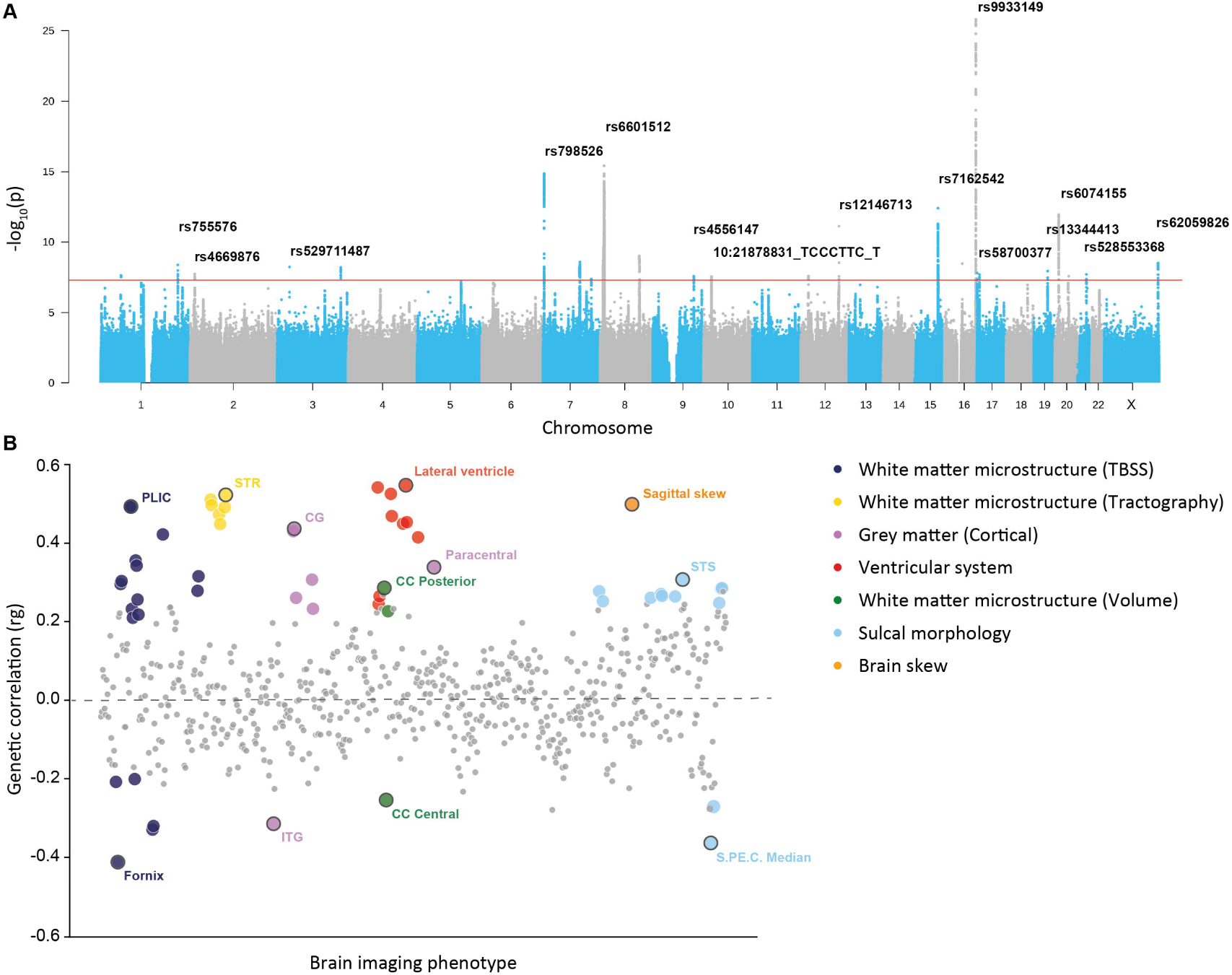
Genetics of interindividual differences in endocranial globularity in a large cohort of present-day humans. **(A)** Manhattan plot of loci associated with interindividual variability in endocranial globularity (N = 33,951); the red line denotes genome-wide significance (p < 5 × 10^−8^). Per chromosome the respective top genome-wide significant loci are annotated with their lead SNP rsID. (**B)** Genetic correlations for endocranial globularity with 667 anatomical brain imaging phenotypes. Large coloured dots denote significant associations after Bonferroni correction, with top hits annotated per imaging category. CG = cingulate gyrus, ITG = inferior temporal gyrus, CC = corpus callosum, PLIC = posterior limb of the internal capsule, S.PE.C Median = median precentral sulcus, STR = superior thalamic radiation, STS = superior temporal sulcus, TBSS = tract-based spatial statistics.

The locus showing the most significant association (lead SNP rs9933149, p = 1.58 x 10^-26^) was located in an intron of *C16orf95* on chromosome 16 (Supplementary Fig. 3). Previous studies have associated this locus with individual variation in an array of neuroanatomical phenotypes ranging from sulcal morphology to white-matter microstructure and ventricular volume^14–24^. Variants in the region have shown association with congenital heart defects^25^ while multigene deletions that encompass the coding region of *C16orf95* have been reported in cases of microcephaly and neurodevelopmental disorders^26,27^.

Most of the significantly associated variants showed small effect sizes, consistent with expectations for a highly polygenic trait and in line with findings from other neuroimaging-based traits28,29. One notable exception was rs529711487, a rare variant (minor allele frequency = 0.002; European 1000 Genomes Phase 3 Reference Panel) on chromosome 3, which had a strong effect on endocranial globularity with a Beta estimate of 6.68 (SE = 1.15) (Supplementary Fig. 4A). The rs529711487 variant was in Hardy-Weinberg equilibrium (pexact test = 1) and had a high imputation quality score (0.91). For this SNP, heterozygous carriers (N = 114) show larger globularity scores compared to peers who are homozygous for the major allele (Supplementary Fig. 4B); since there are no homozygous carriers of the minor allele in the UK Biobank neuroimaging cohort, it was not possible to test whether it acts in an additive or dominant manner.

Introgressed Neandertal alleles on chromosomes 1 and 18 that were associated with endocranial globularity in a prior targeted study of smaller cohorts^5^ exhibited the same direction of effect towards reduced globularity in UK Biobank (rs28445963, beta (SE) = -0.19 (0.22), p = 0.37; rs72931809, beta (SE) = -0.16 (0.21), p = 0.45) but associations were not significant.

Next, we used the aggregated GWAS signal across the entire genome to gain further insights into the underlying biology. Estimates of SNP-heritability indicated that the captured genetic variation accounted for 32.6% (SE = 3.4%) of the observed phenotypic variation in endocranial globularity in the cohort. As genetic sex has distinct influences on brain (development), it is generally considered a possible confound in GWASs. To supplement the phenotypic analysis we also performed sex-stratified GWASs, finding a high genetic correlation between the male (N = 16,148) and female (N = 17,803) GWAS (rg (SE) = 0.9363 (0.0676)), p = 1.245 x 10^-43^) (Supplementary Fig. 5). Notably, with the exception of one indel (8:108402192_AGC_A, male GWAS), all independent significantly associated variants in sex-stratified GWASs were also identified as genomic regions associated with endocranial globularity (independent lead SNPs and SNPs in LD (r2 >0.6)) in the primary GWAS (Supplementary Table 2). Thus, endocranial globularity shows no pronounced sexual dimorphism, either phenotypically or genetically.

We went on to study relationships between endocranial globularity and a selection of other relevant phenotypes, by estimating genetic correlations to measure the extent to which shared genetic variation contributes to shared phenotypic variability across traits.

We used brain imaging derived phenotypes available in UK Biobank to assess whether certain anatomical regions or specific metrics show significant genetic overlap with interindividual variation of endocranial globularity, encompassing cortical and subcortical estimates as well as white-matter microstructure estimates derived from tract-based spatial statistics (TBSS) and probabilistic tractography. Further, we included metrics detailing sulcal brain morphology^14^ and brain skew^30^, traits with potential relevance to human evolution^31–34^ that are not part of the standard UK Biobank processing pipeline. Of the 667 neuroanatomical phenotypes that we analysed, 56 showed significant genetic correlations with endocranial globularity (α_Bonferroni_ = 0.05 / 667 = 7.5x10^-5^) (Fig. 2B, Supplementary Table 3), with strongest correlations for the ventricular system, distinct white-matter tracts, certain sulcal width measures, and sagittal brain skew. Both hemispheres contributed similarly to these correlations, while cortical (surface area, thickness) and subcortical (volume) measures generally showed only small genetic overlaps with endocranial globularity.

Because facial form was recently proposed to be a main driver of endocranial shape variation during hominin evolution^7^ we tested whether any of the identified genomic regions associated with endocranial globularity (independent lead SNPs and SNPs in LD (r^2^ >0.6)) were also associated with individual differences in facial morphology in previous GWAS efforts^35–51^. This analysis highlighted four independent signals from four different genomic loci which showed both an association here with endocranial globularity and in previous facial shape studies, mostly with the morphology of the nose (Supplementary Table. 4).

Some have proposed links between the unique globularity of the human brain and the evolution of cognitive capacities and emergence of complex human traits^52^. At the same time, evidence from a range of sources supports a reappraisal of Neandertal capacities for complex behaviour, suggesting that our archaic cousins had more cognitive sophistication than previously appreciated^53–56^. We tested for shared genetic underpinnings of endocranial globularity with a curated set of complex human traits across several biological domains where an evolutionary relevance was previously highlighted. Our choice was motivated by prior studies which highlighted the phenotypic legacy of previous admixture events over trait domains linked to metabolism, dermatology, the cardiovascular and gastrointestinal system, cognition, and skeletal features, as well as neuropsychiatric disorders^4,57–67^ (Fig.3).

**Fig. 3.**
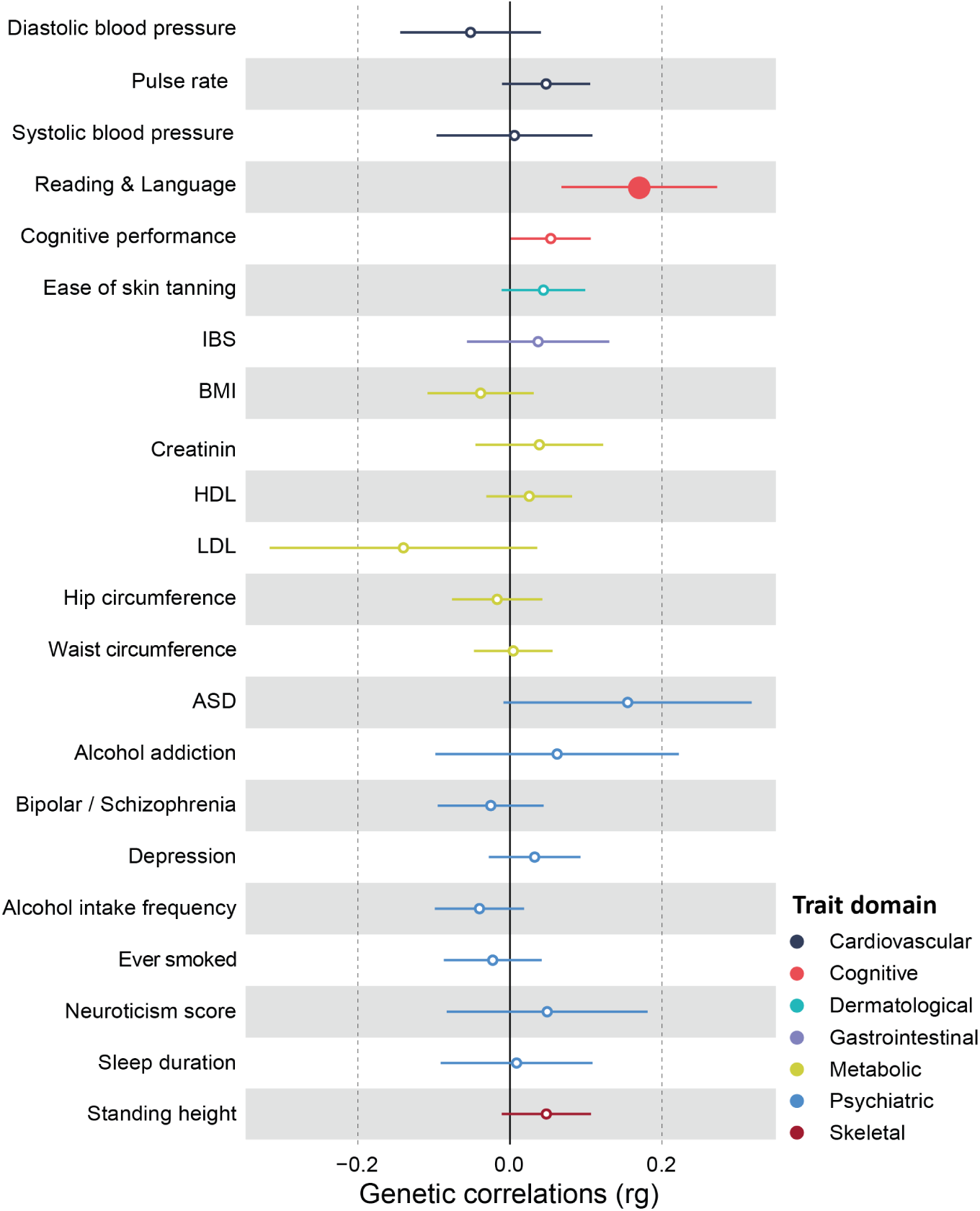
Genetic correlations between endocranial globularity and disease phenotypes and other complex traits. Large coloured dots denote significant associations after Bonferroni correction for the 22 phenotypes that were studied. Genetic correlation (rg) is presented as a dot, and error bars indicate the standard error. ASD = Autism spectrum disorder, BMI = Body mass index, HDL = high-density lipoprotein, IBS = Irritable bowel syndrome, LDL = low-density lipoprotein;

A multivariate GWAS on reading- and language-related traits^68^ was the only one to demonstrate a significant genetic correlation (rg(SE) = 0.1704 (0.0523), p = 0.0011; α_Bonferroni_ = 0.05 / 22 = 0.0023). In contrast, a measure of overall cognitive performance, as well as all other trait domains assessed here, exhibited no significant genetic overlap with endocranial globularity (Supplementary Table 5).

## Functional enrichment in gene expression and chromatin data

To gain insights into functions of genes related to endocranial globularity, we first examined gene-based associations using MAGMA’s gene-based test as implemented in FUMA^69^. 77 genes were significantly associated with interindividual variation in endocranial globularity (α_Bonferroni_ = 0.05 / 19,312 = 2.59 x 10^-6^; Supplementary Table 6) with the top signal seen for *C16orf95*, in line with the SNP-based associations. We then performed MAGMA gene-property analysis to test for positive associations between tissue-specific gene expression profiles and gene-based p-values from the GWAS. Using two gene expression datasets (GTEx, Brainspan) (α_Bonferroni_ = 0.05 / (54 + 11) = 7.69 × 10^-4^) we identified significant enrichment in adult tissue (GTEx)^70^ highlighting the tibial nerve as well as several non-brain tissues, mainly attributed to the cardiovascular, gastrointestinal (colon and oesophagus) and female reproductive systems (cervix, uterus, fallopian tube) (Fig. 4A, Supplementary Table 7). Further, enrichment in brain tissues sampled through development (Brainspan)^71^ highlighted late-prenatal and early childhood brain tissues (Fig. 4B, Supplementary Table 7).

**Fig. 4.**
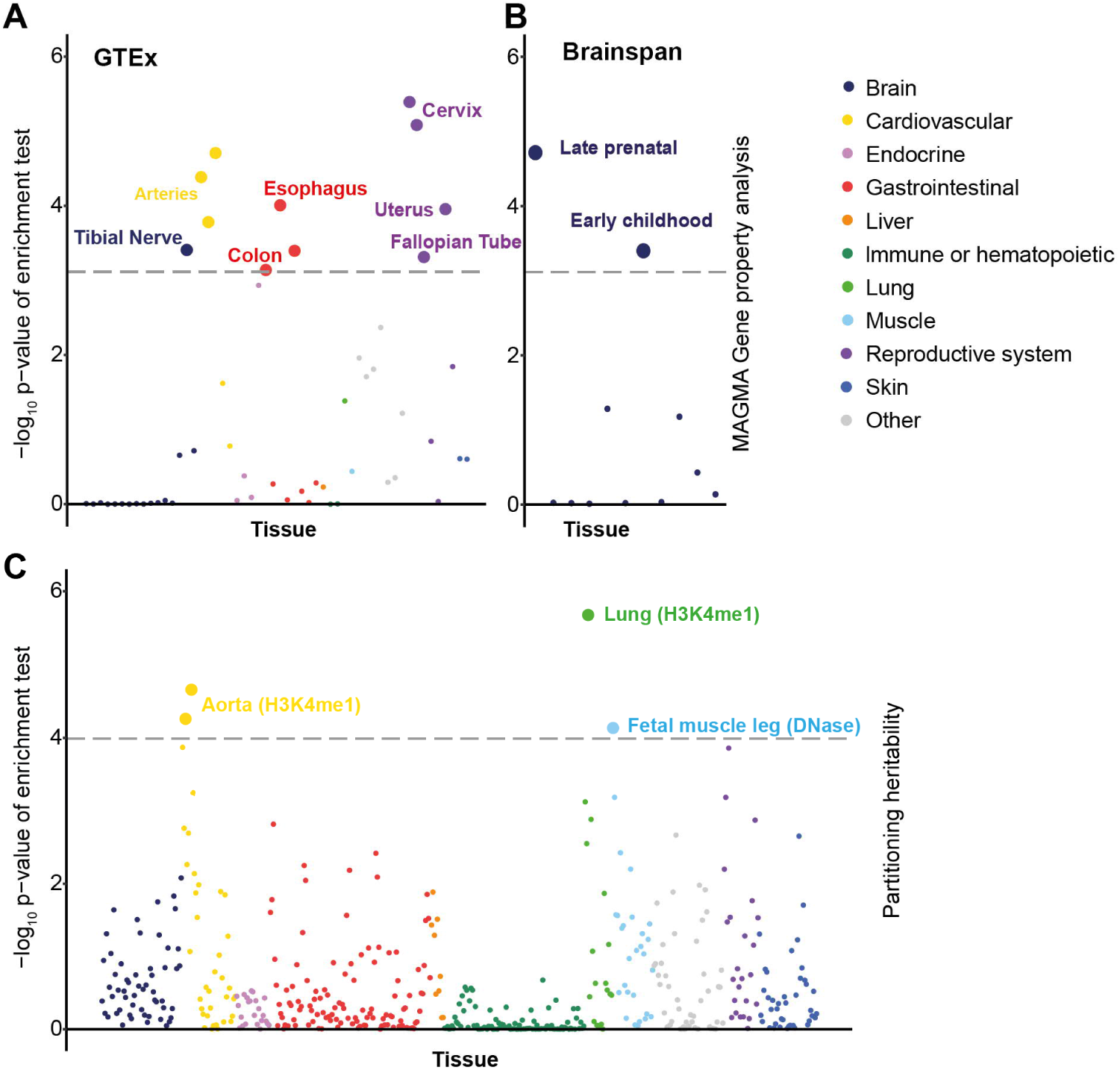
Analyses of expression profiles in adult and brain developmental tissue as well as tissue-specific chromatin signatures. Enrichment -log_10_ p-values of MAGMA gene-property analysis in gene expression data from (A) adult tissue and (B) brain tissue from different developmental time points. (C) Partitioned heritability enrichment - log_10_ p-values in active regulatory elements across tissues and cell types. In all panels, the dashed line denotes the p-value threshold for significant enrichment after Bonferroni correction.

We next tested for (brain) cell type enrichments using two single-cell RNA-sequencing datasets covering foetal and adult brain tissue which highlighted significant enrichments in endothelial cells of adult tissue (Supplementary Fig. 6, Supplementary Table 8).

To complement the gene expression analyses, we investigated enrichment of globularity associations in tissue-specific chromatin signatures, using LDSC partitioned heritability analysis^72,73^. We found significant enrichment (α_Bonferroni_ = 0.05 / 489 = 1.02 x 10^-4^) in regions with DNase hypersensitivity, a marker for active chromatin, in foetal skeletal muscle (Fig. 4C, Supplementary Table 9) and in regions with histone-3 lysine-4 monomethylation (H3K4me1), a marker for enhancer regions, which again highlighted the cardiovascular system but also showed enrichment in lung tissue (Fig. 4C, Supplementary Table 9).

## Partitioned heritability analysis with evolutionary annotations of the genome

Focusing on the timescale over which endocranial globularity is thought to have emerged on the *Homo sapiens* lineage (i.e., within ∼300kya, see Figure 1A), we investigated relationships between our association data and genomic annotations that tag relevant evolutionary events.

First, we used LDSC partitioned heritability analysis^72,73^ to test the contributions of archaic admixture to the overall SNP-based heritability of the trait. Here, we analysed (i) Neandertal introgressed alleles resulting from admixture events ∼50-60k years ago^74^ and (ii) archaic deserts, which represent long stretches in the human genome that have a significant underrepresentation of introgressed alleles^75^. This analysis revealed a significant (α_Bonferroni_ = 0.05 / 2 = 0.025) heritability depletion in archaic deserts (enrichment = 0.61, SE = 0.14, p = 0.0115), indicating that common DNA variants in these regions explain a lower proportion of the total SNP-based heritability of endocranial globularity than expected under complete polygenicity (Supplementary Table 10).

Next, we investigated links between endocranial globularity and two further sets of evolutionary annotations that match the relevant timescale: (i) ancient selective sweeps, as defined by extended lineage sorting^76^, and (ii) anatomically modern human-derived differentially methylated regions (AMH-derived DMRs) identified using skeletal DNA methylation maps of Neandertals, Denisovans, and present-day humans^77^. Since these annotations cannot be tested reliably with existing LDSC heritability partitioning tools^78^, we adopted a MAGMA gene-set approach^79^ using two custom gene-sets covering genes that fall within ±1kb of the relevant evolutionary annotations. We found that common variants associated with endocranial globularity are significantly (α_Bonferroni_ = 0.05 / 2 = 0.025) enriched in and near genes (±1 kb) co-located with ancient selective sweeps (Beta = 0.163, SE = 0.061, p = 0.004), but not in those co-located with AMH-derived DMRs (Beta = 0.02, SE = 0.003, p = 0.333) (Supplementary Table 11).

## Discussion

Our results point to proximate developmental, and evolutionary causes underlying a uniquely modern human anatomical phenotype. The origins of endocranial globularity have been much debated, with some studies emphasizing roles of genes related to neurogenesis and myelination^2,5^, while others propose that endocranial shape changes in *Homo sapiens* are driven by facial gracilization linked to dietary and lifestyle differences^7^. In the current study, we approached this longstanding question by investigating interindividual differences in endocranial globularity in nearly 34,000 present-day living participants, identifying 28 loci and 77 genes associated with the trait, and revealing a SNP-heritability comparable to (and in some cases larger than) other neuroanatomical traits which have been studied via brain imaging genomics^15,21,80^.

We found that genetic variation associated with endocranial globularity is shared with that of several neuroanatomical metrics covering a variety of brain regions, highlighting the multifactorial nature of this evolutionarily-defined phenotype. Particularly prominent here is the shared genetic make-up with the ventricular system, raising the possibility that rounder endocrania may in part be due to increased hydrostatic pressure of cerebrospinal fluid during development. Moreover, these results go beyond confirming prior hypotheses concerning a role for white-matter changes^2,5^, by revealing which white-matter tracts show the strongest genetic overlaps with endocranial globularity (Supplementary Table 3).

While the globularization of the brain is thought to have contributed to the emergence of complex human-specific traits^52,53,81,82^, adaptive evolution might have increased our susceptibility to certain (brain-related) disorders^57,58,61^. In the present study, we tested 22 phenotypes in a range of categories where prior studies have suggested an impact of previous admixture events. Across all trait domains we found at most very subtle genetic relationships with endocranial globularity, and only one of the traits that we tested met statistical significance. This is in line with recent studies finding that Neandertal variants overall show little association with diseases seen in present-day humans, specifically with psychiatric and neurological disorders^58,63^. More generally, our findings indicate that the trend towards a more globular brain is not per se linked to increased disease susceptibility and seems fairly independent of other trait domains previously linked to Neandertal introgression. While we did not expect straightforward correlations between brain shape and complex human traits, the absence of shared genetic architecture of endocranial globularity with traits previously linked to Neandertal introgression might in part also reflect the focus on common variants in this study, given that a large percentage of Neandertal introgressed fragments is rare with allele frequencies below 1%^83^.

Reading- and language-related skills were the only cognitive traits that showed a significant genetic correlation with endocranial globularity. However, this does not necessarily imply the absence of language in archaic hominins. As our analysis revealed only a marginal, non-significant genetic overlap with general cognitive abilities, this finding may rather indicate a continuous refinement of language-related abilities in the *Homo sapiens* lineage after the split from Neandertals. Indeed, based on data from an array of palaeoanthropological sources, it has been suggested that Neandertals had more complex communication abilities than previously appreciated, potentially reflecting a deeper evolutionary history for speech and language capacities^84–87^. This is consistent with archaeological evidence of complex and sophisticated Neandertal behaviours, suggesting cognitive and behavioural similarities between Neandertals and contemporary fossil *Homo sapiens*^53–56^. Given both the shared and diverging genetic architecture with anatomical brain traits, subtle differences in brain circuitry which might emerge during the globularization phase of brain development^2,5,6^ may have refined key brain structures important for language circuitry that has subsequently also been recruited for processes involved in reading. Intriguingly, anatomical measures of the superior temporal sulcus, a key brain region involved in linguistic processing, show high genetic correlations with both endocranial globularity and reading-/language-related traits^68^.

Focusing on functional implications of identified associations, gene-property analysis highlighted an enrichment of variants associated with endocranial globularity in genes expressed in the brain during prenatal development and in early childhood. This finding links directly to prior work on skulls which places the ontogenetic emergence of this phenotype within early development^5,6,88,89^.

Several lines of evidence point to an association between endocranial globularity, metabolic supply and the circulatory system. First, endocranial globularity showed strong genetic correlations with the ventricular system, including the choroid plexus. Second, via gene-property analysis, we found that genes associated with this trait have enriched expression in tissues linked to the cardiovascular system as well as in adult brain endothelial cells, a cell type also linked to the brain’s metabolite supply^90^. Third, using heritability partitioning, we observed enrichment of heritability within regulatory elements linked to the cardiovascular system. In addition to the cardiovascular system, genes associated with endocranial globularity also showed significant enrichment in female reproductive tissues. The association of endocranial globularity and the cardiovascular system highlights the advantages of our approach to uncover novel associations which open up unexpected avenues for further research. This is consistent with recent research that has shown strong phenotypic and genetic links between the heart and the brain^91^ and even uncovered possibly genetic causal links between cardiovascular endophenotypes and neurological health.

While Gunz & Tilot et al.^5^, using a smaller cohort, identified a number of Neandertal introgressed alleles associated with elongated endocranial shape in present-day humans, our genome-wide approach revealed a broader signal of SNP-heritability depletion within archaic deserts. This may suggest that archaic deserts lack the genetic loci and regulatory elements that contribute to the formation of the globular shape of the human endocranium during early development, similar to earlier findings^78^ showing archaic desert heritability depletion for cortical surface area. Further, the gene-set analysis indicated a significant enrichment of globularity-associated variants within genes that are co-located with ancient selective sweeps. Thus, genes that are co-located with haplotypes fixed on the human lineage after the split from Neandertals and Denisovans (∼450-750 kya)^13^, but before the split of modern human populations (∼100-120 kya)^66^, may have played an important role in the evolution of human endocranial globularity. The implicated time period partially overlaps with that observed from the fossil data, which suggests endocranial globularity emerged in the *Homo sapiens* lineage over the last ∼300 kya. Overall, combining fossil and genetic data narrows down the presumable time window for the evolutionary emergence of endocranial globularity to the last ∼300-120 kya.

An inherent limitation of the GWAS approach is that it cannot directly assess the impact of genetic changes that have become fixed over the course of hominin evolution, as the basis of the analysis relies on both genetic and phenotypic interindividual variation present in the cohort under study. In addition, for pragmatic reasons, our study focused on individuals of European ancestry, highlighting the need for further research of well-powered neuroimaging genomics cohorts from diverse population backgrounds (as they become available) to gain a comprehensive understanding of the biological mechanisms underlying aspects of human neuroanatomy. It is important to keep in mind ethical implications of research on fundamental human traits; Box 1 outlines key issues, highlighting the scope and limitations of GWAS and providing guidelines for interpreting such genetic findings.

**Box 1 General ethical considerations and implications linked to brain shape and its genetic underpinnings**

Our study leveraged the interindividual differences in an evolutionarily derived phenotype unique to present-day humans through analyses of large-scale neuroimaging genetics, to gain novel insights into biological processes underlying human variation and evolution. We uncovered unexpected signals that lead to new, testable hypotheses for future work when more fine-grained datasets become available. Using neuroimaging genomic data spanning perinatal and infant development will then allow for more detailed insights into the ontogeny of endocranial shape while the latest data from single-cell transcriptomics and epigenomics will help to further disentangle the involvement of metabolic supply, the circulatory and female reproductive system. Further, availability of large-scale exome and whole-genome sequencing data will shed more light on the contribution of rarer variants with larger effect sizes, whose functional implications and interplay with more common variants are not yet clear.

Taken together, these findings suggest that human endocranial shape and brain function may have co-evolved with both the circulatory and reproductive systems. In such a scenario, selection acting on one system would also affect the other; thus, changes affecting, e.g., metabolite supply, energy demand, pregnancy, or fertility would also affect brain structure and endocranial shape, possibly as indirect by-products.

## Historical context of craniology and phrenology warrants a careful and vigilant ethical approach

Endocranial imprints can provide direct empirical evidence of brain evolution revealing evolutionary changes of brain size, shape and sulcal morphology in palaeoanthropology^2,6,92–96^. However, given the historical burden of craniometry, using endocranial metrics as phenotypes and correlating them with anatomical or behavioural traits requires vigilant and thorough ethical considerations to avoid misappropriation of presented results^97,98^. Craniometry, the study of skull shape, size, and proportions, was prevalent in anthropology during the 18th and early 19th centuries. It misused measurements, like the cervical index (the ratio of a skull’s maximum width to its length), to wrongly suggest racial differences in intellectual capabilities. These ideas were absorbed into phrenology, which claimed to predict personality, mental health, and criminal tendencies based on skull measurements, providing a pseudo-scientific basis for racist beliefs, slavery, colonialism, and later, 20th-century eugenics^99,100^. Despite being discredited as pseudoscience^101,102^, the remnants of racial craniology and phrenology still influence contemporary studies linking brain and behaviour^103^.

## Brain shape as a window into our evolutionary past

In this study we make use of the latest advances in palaeoanthropology and endocranial shape analysis from Neandertal fossils combined with brain imaging data from thousands of present-day living humans to create a singular metric for endocranial globularity^5^. As the globular shape of the human braincase represents one of the most distinct anatomical differences between *Homo sapiens* and our closest living and extinct relatives, endocranial globularity functions here as an evolutionary phenotype^2,6^. Investigating associated genetic factors underlying this evolutionary phenotype offers a novel approach to study the more elusive aspects of brain evolution and might lead to new insights into differences and similarities compared to our closest extinct relatives, with no intent or scientific grounds to draw any conclusions for an individual based on differences in their endocranial globularity.

## What can genetics tell us about human endocranial globularity today?

Genome-wide association analyses of endocranial globularity highlight the polygenicity of this trait; that is, there are thousands, if not tens of thousands, of DNA variants acting together to shape this particular aspect of our anatomy. While identified associations do not inherently imply causal relationships, they can open up new avenues of scientific inquiry for human evolution^104,105^. Overall, endocranial globularity showed a SNP-heritability (the proportion of interindividual variation that may be accounted for by common genetic variants) of 30%, comparable to other brain morphological traits^80,106,107^. Still, this means that only a third of the interindividual variation in endocranial globularity can currently be explained by factors at the genetic level. While larger sample sizes and a focus on rare variants might increase the genetic contribution, this level of heritability indicates that a substantial proportion of variability in this trait is still unexplained and can be attributed to non-genetic factors such as environmental influences^104^ and developmental stochasticity. Even though each identified genetic variant associated with endocranial globularity shows only minor effects, the aggregated signal was correlated with a range of brain morphological traits, while most of the health and complex human traits that we studied did not show significant genetic correlation. Overall, these analyses can reveal the aetiological heterogeneity of endocranial globularity and other complex traits, and help us to better understand their shared biology. Yet, it is important to note that genetic correlations do not imply causal links between phenotypes^108^.

We emphasize that our results and summary statistics do not provide a scientific foundation for deterministic predictions of human endocranial shape. The findings represent statistical trends within the studied population and cannot be applied to individuals. Additionally, given that different populations show differences in allele frequencies and patterns of linkage disequilibrium (i.e. co-inheritance of neighbouring DNA variants that lie close to each other on the same chromosome), this also means that the results should not be taken out of context to make any inferences about other ancestry groups or to draw comparisons between populations.

## Methods

### Dataset

For lifespan analyses we included 644 cognitively healthy adults (age range, 18 – 88 years) obtained from the Cambridge Centre for Aging and Neuroscience (CamCAN) repository (available at http://www.mrc-cbu.cam.ac.uk/datasets/camcan/)^9,10^. Ethical approval was obtained from the Cambridgeshire Research Ethics Committee and participants gave written informed consent.

All data for the described neuroimaging genetics analysis were obtained from the UK Biobank (UKB) under the research application 16066 with Clyde Francks as the principal investigator. Detailed descriptions of the data used as well as sample-specific pre-processing and quality control (QC) are described below. The UK Biobank received ethical approval from the National Research Ethics Service Committee North West-Haydock (reference 11/NW/0382), and all of their procedures were performed in accordance with the World Medical Association. Informed consent was obtained for all participants by UK Biobank with details about data collection and ethical procedures described elsewhere^11,109^.

### Endocast segmentation and measurement protocol

Based on computed tomographic (CT) scans of recent and fossil crania, we used the endocranial shape differences between present-day humans (N = 89) representing world-wide cranial shape variation^2^, and fossil humans (N = 20) as a framework for evolutionary shape changes^5^. The fossil sample comprises eight *Homo erectus* s.l. (KNM-ER 3733, KNM-ER 3883, KNM-WT 15000, OH 9, Ngandong 14, Ngawi, Sambungmacan 3, Sangiran 2), ten *Homo neanderthalensis* (Amud 1, Feldhofer 1, Gibraltar 1, Guattari 1, La Chapelle aux Saints, La Ferrassie 1, Spy 1, Spy 2, Saccopastore 1, Le Moustier 1), and two *Homo heidelbergensis* s.l. crania (Kabwe, Petralona). We first used semi-automated segmentation to create virtual endocranial imprints in the software Avizo 3D (Version 9; FEI SAS, Thermo Fisher Scientific) following protocols detailed in Neubauer et al.^2,88^. We also segmented the endocranial surface of the MNI 152 template, an average of 152 normal MRI brain scans. On each endocranial surface we then digitized a mesh of 935 anatomical landmarks, curve semilandmarks^110^ and surface semilandmarks following the protocol described in Neubauer et al.^2^To establish geometric correspondence of the semilandmarks across individuals we allowed curve semilandmarks to slide along tangents to the curves, and surface semilandmarks to slide on tangent planes to the surface, minimizing the thin-plate spline bending energy following Gunz et al.^110^. Missing landmarks or semilandmarks were estimated based on minimal bending energy during this sliding step^111,112^.

### Sample level quality control and pre-processing of the neuroimaging data

For the lifespan analyses we used available T1-weighted images within the CamCAN data set. All details regarding the MRI acquisition are described in detail in Taylor et al.^9^. All available T1-weighted images were further pre-processed using *fsl_anat* as part of the FSL toolbox (v 5.1.0) to reorient the images to MNI standard orientation, automatically crop and bias-correct the images, and apply an initial brain extraction which was followed by a secondary brain extraction using FSL *bet*. Images were subsequently aligned to the MNI-T1-1mm standard template using FSL *flirt* to derive the inverse affine transformation matrix for each of 644 individuals.

For UKB data we first applied genetic sample level quality control measures to derive a final study sample that would also be suitable for further genetic analysis. As the first step, participants with available imaging data (N = 40,681, release 2020) were extracted from the full UKB imputed genotype dataset^11^. Subject-level QC parameters were then applied to the 39,678 individuals for whom both neuroimaging and genotype data were available. This involved excluding individuals with a mismatch of their self-reported (UKB data field 31) and genetically inferred sex (UKB data field 22001), as well as individuals with putative aneuploidies (UKB data field 22019), or individuals who were determined as outliers based on heterozygosity (PC corrected heterozygosity >0.1903) or genotype missingness rate (missing rate >0.05) (UKB data field 22027)^11^. For all participants with self-reported ‘White European’ ancestry (UKB data field 21000) who passed the above described sample level quality control, a Bayesian outlier detection algorithm (lambda = 40) was applied (*abberant*^113^). This algorithm identified dense clusters of participants with similar genetic ancestry along PC1-PC2, PC3-PC4 and PC5-PC6, where only participants at the intersection of all three principal component clusters were subsequently included. Lastly, we identified pairs of individuals with a kinship coefficient > 0.0442 (UKB data field 22021^11^) and available imaging data, and excluded one individual from each pair, prioritizing the exclusion of individuals related to a larger number of other individuals to maximize the overall sample.

This process resulted in 34,861 participants passing the described genetic sample level QC, of which 33,996 also had usable imaging data (T1 folder not declared ‘unusable’). From this sample, we then made use of imaging-derived phenotypes and pre-processed image data generated by an imaging-processing pipeline developed and run on behalf of the UK Biobank. All details regarding image acquisition and subsequent applied processing pipelines are available on the UK Biobank website (http://biobank.ctsu.ox.ac.uk/crystal/refer.cgi?id=2367), and in the respective brain imaging documentation (http://biobank.ctsu.ox.ac.uk/crystal/refer.cgi?id=1977), as described elsewhere^114,115^. We used the provided linear affine transformation matrices of each individual, which was part of the standard T1-weighted MRI processing pipeline (UKB data field 20252; T1_to_MNI_linear.mat). We further used FSL (v5.1.0, *convert_xfm*) to derive the inverse affine transformation matrix for each of the 33,996 participants.

### Globularity scores derived from structural MRI scans and quality control

To quantify the endocranial shape differences between modern humans and Neandertals, we combined geometric morphometrics^8^ using scripts in Mathematica (Version 12; Wolfram Inc.) with standard neuroimaging data. In Mathematica, we applied the derived inverse affine transformation matrices for each individual from both datasets (CamCAN, UKB) to the 3D coordinates of the dense mesh of 935 landmarks and semilandmarks on the MNI 152 template, thereby bringing these coordinates into the native anatomical space of each individual. Using Procrustes superimposition^116^ we standardized position, orientation, and scale. From these Procrustes shape coordinates we then computed the difference between the mean endocranial shape of Neandertals, and the mean endocranial shape of present-day humans. To calculate the globularity scores, we projected each individual onto that multivariate vector, and multiplied the resulting scalars by 1000. In this summary metric, more elongated (Neandertal-like) endocrania have low values, and more globular endocrania have high values.

### Quality control of the derived phenotype

For both datasets (CamCAN, UKB) we manually checked the respective T1-weighted image of any individuals presenting with extreme globularity scores (Q1/Q3 +/-1.5 IQR) for crude deviations in brain shape from derived globularity scores. Further, the created inverse transformation matrix for all outliers was applied to the MNI 152 standard template using FSL *flirt* and the derived image overlaid on each individual’s initial T1-weighted image, respectively. All overlays were manually inspected and individuals with crude misalignments (N = 45) excluded from subsequent analysis, resulting in 644 and 33,951 individuals for CamCAN and UKB, respectively.

### Characterization of phenotype

Previous work suggested a subtle relationship between age and globularity scores^5^. We refined this characterization and systematically assessed the relationship between age and endocranial globularity during adulthood within the CamCAN lifespan dataset (N = 644) as well as the UKB dataset (N = 33,951), also accounting for possible interactions with sex. To do so, we used a univariate linear regression implemented in R [*lm*], including both age at time of scanning (UKB data field 21003; instance 2.0) and age^2^ terms and sex (UKB data field 31) as covariates, as well as possible interaction term (sex*age, sex*age^2^) in our model. For a more detailed description of possible sex dimorphism, we assessed the effect of sex on endocranial globularity using a univariate linear regression, where sex was included as the main binary predictor (N_Male_ =16,148; N_Female_ = 17,803), while controlling for age. We extracted the t-statistic for the ‘sex’ term to calculate Cohen’s d effect sizes and 95% confidence intervals using the psych package (version 2.1.9) in R^117^.

Further, we investigated relationships with estimated total intracranial volume (UKB data field 26521, instance 2.0, N = 33,577), and subject-specific health data (UKB Category 100074 and 2002). As before, we applied a univariate linear regression to assess relationships between brain size and endocranial globularity, using both age and sex as covariates. For subject-specific health data, we screened provided health information for distinct neuroanatomical and neurodegenerative disorders that might potentially have an influence on human brain shape, combining both supplied ICD9/10 information (UKB data fields 41202, 41203, 41204, 42205) and self-reported data (UKB data field 200002) which were collected in a verbal interview at the assessment centre. In each case, all instances up to the imaging visit (instance 2) were screened. This screen highlighted 27 participants with an ICD9 diagnosis and 1022 individuals with an ICD10 diagnosis, while 1808 individuals indicated a neuroanatomical or neurodegenerative diagnosis in the self-reported health screen. As some participants had multiple records, this led to a total of 2,338 unique diagnosed participants while the rest of the cohort (N = 31,613) were treated as control participants. We assessed the relationship with these disorders using a univariate linear regression, where diagnosis (case-control status) was included as the main binary predictor, while controlling for sex and age. We extracted the t-statistic for the ‘diagnosis’ term to calculate Cohen’s *d* effect sizes and 95% confidence intervals using the *psych* package (version 2.1.9) in R^117^.

### Genome-wide association analysis

For all individuals in the UKB sample (N = 33,951), imputed single-nucleotide polymorphism (SNP) genotype data (UKB Category 263, bgen files; imputed data v3-released March 2018) were extracted and variant level QC and SNP statistics were computed using qctool v2.0.2 (https://www.well.ox.ac.uk/∼gav/qctool_v2/). We excluded variants with a minor allele frequency below 0.001, Hardy-Weinberg equilibrium test p-value < 1×10^-6^, imputation quality INFO score <0.7 (included in the imputed UKB data) and multi-allelic SNPs. As recommended, for the X chromosome only Hardy-Weinberg equilibrium p-values based on the female dataset were checked^118^. This procedure resulted in 15,030,319 SNPs, of which 499,719 were located on the X chromosome.

A linear regression model as implemented in PLINK 2.0 (www.cog-genomics.org/plink/2.0/)^119,120^ was used to test for the association of endocranial globularity with sample specific dosages, using an additive model. Here, PLINK converted the raw probabilities stored in the .bgen files to dosages. Covariates were scaled to have unit variance, by using the *--covar-variance-standardiz* flag. To account for dosage compensation (DC) due to X chromosome inactivation in females, we assumed full DC and treated males as homozygous females with genotypes coded as (0,2) which represents the default model in PLINK 2.0^121^. We included age (UKB data field 21003-2.0), age^2^, sex (UKB data field 31-0.0), sex-by-age and age^2^ interactions, the first 10 genetic principal components (UKB data fields 22009-0.1 to 22009-0.10), genotype measurement array (UKB data field 22000-0.0), and dummy variables for assessment centre (UKB data field 54-2.0), as well as several imaging-related covariates, namely scanner position parameters (X, Y and Z position: UKB data fields 25756-2.0, 25757-2.0 and 25758-2.0), T1 signal-to-noise ratio (UKB data field 25734-2.0), and T1-contrast-to-noise ratio (UKB data field 25735-2.0) as co-factors.

For sex-stratified analysis we used the *--keep-male* or *--keep-female* flags as implemented in PLINK 2.0, which decreased the overall sample size to 16,148 and 17,803 individuals, respectively. The sex-stratified association tests were not adjusted for sex or any interaction terms thereof. The Manhattan plot was generated using the “qqman” R function^122^ (qqman package v0.1.8), while QQ plots were added with a custom function (https://genome.sph.umich.edu/wiki/Code_Sample:_Generating_QQ_Plots_in_R by Matthey Flickinger). For sex-stratified GWAS, the Miami plot was created with the custom R function gmirror (https://github.com/RitchieLab/utility/blob/master/personal/ana/hudson-paper/hudson-paper-figures-code.R).

Subsequently, LD score regression (v1.0.0)^123^ was used to estimate SNP heritability for endocranial globularity and to calculate genetic correlation estimates between the male and female GWAS. For all these analysis streams and any following analysis using *ldsc*, only autosomes were included, since the X chromosome is not supported by the *ldsc* software.

### Genetic correlations with brain imaging phenotypes

Publicly available GWAS summary statistics of brain imaging derived neuroanatomical traits were obtained via the Oxford Brain Imaging Genetics Server^15^ (http://big.stats.ox.ac.uk/). Brain imaging derived phenotypes (IDPs) encompassed surface T1 structural phenotypes (UKB Category 110), namely regional grey matter volumes (FAST, UKB Category 1101), subcortical volumes (FIRST, UKB Category 1102), Freesurfer ASEG (UKB Category 190) and Freesurfer DKT (UKB Category 196), as well as diffusion imaging phenotypes (UKB Category 107), where we selected both fractional anisotropy (FA) and mean diffusivity (MD) from the diffusion MRI skeleton measurements (UKB Category 134) and diffusion weighted means measures (UKB Category 135), respectively. We also included summary statistics for two further brain IDPs, namely sulcal morphology measures^14^ and brain skew^30^. This resulted in 1017 traits, where only those with a SNP-heritability larger than 0.1 were used, to avoid signal inflation, leading to a total of 667 final brain IDPs. A description of all IDPs used in this study can be found in Supplementary Table 3. LD score regression^123^ was used to assess genetic correlations between endocranial globularity and the brain IDPs, with Bonferroni correction to adjust for multiple testing of 667 phenotypes.

As GWAS statistics for facial morphology traits were mostly available from multivariate screens and hence lacked beta estimates, LD score regression was not applicable to test for possible genetic correlation. We therefore used a lookup approach as an alternative, to determine if any of the genome-wide significant loci associated with facial morphology (as reported in Supplementary Tables 1 and 3 of White et al.^35^) overlapped with identified genomic regions associated with endocranial globularity (independent lead SNPs and SNPs in LD (r^2^ > 0.6)).

### Genetic correlation analysis with complex human traits

Several studies have highlighted potential relevance to human evolution for certain brain-related disorders including depression, autism, schizophrenia, and addiction^57,60,61,124^, inflammatory conditions^64^, as well as cognitive traits, with language being the most prominent^66,86,87^. Further, the legacy of previous admixture events was shown to influence trait variability in several categories, ranging from metabolism to dermatological traits, as well as the cardiovascular and gastrointestinal system, and skeletal form. We curated a list of 22 traits within these domains where high quality GWAS summary statistics were available and used LD score regression^123^ to estimate genetic correlations between endocranial globularity and these traits. We made use of publicly available GWAS summary statistics for schizophrenia and bipolar disorder^125^, depression^126^, autism spectrum disorder^127^, alcohol dependence^128^, inflammatory bowel disease^129^, cognitive performance^130^, and a multivariate GWAS on reading- and language-related traits^68^. For all other traits, LDSC formatted summary statistics were downloaded from the Neal Lab heritability server^131^ (https://nealelab.github.io/UKBB_ldsc/index.html). Bonferroni correction was used to correct for multiple testing of 22 traits. A detailed overview of results and a description of all traits used in this study can be found in Supplementary Table 5.

### Functional Mapping and Annotation of GWAS analysis results

For functional annotation of GWAS results, we used the SNP2GENE function implemented in FUMA ^69,132^ (Functional Mapping and Annotation of Genome-Wide Association Studies; https://fuma.ctglab.nl/; version v1.3.7) to highlight associated genomic loci and independent significant SNPs (r^2^ > 0.6).

### Gene and gene-property analysis in MAGMA

MAGMA^79^ (version 1.08), integrated in the SNP2GENE function in FUMA, was used for gene-based, gene-set, and gene-property analyses. For gene-based analysis (SNP-wide mean model) the European 1000 Genomes Phase 3 Reference Panel^133^ was used and the gene window was set to 35 kb upstream and 10 kb downstream^134^ to determine gene-based p-values, with other settings using default values. MAGMA controls for gene size, the number of SNPs within a gene, and LD structure, by default.

We tested for enrichment of low gene-based p-values in specific tissue and cell types using MAGMA gene-property analysis, where we selected adult tissue samples from GTEx V8 (http://www.gtexportal.org/home/datasets), as well as gene expression data of developmental brain samples (http://www.brainspan.org). Additionally, we chose two single-cell RNA-sequencing datasets covering fetal (prefrontal cortex, 8-26 weeks post conception; Gene expression omnibus (GEO) accession number GSE104276) and adult (PsychENCODE Adult, http://resource.psychencode.org/) brain tissue for the cell type-specific analysis, also implemented in FUMA^132^. Bonferroni correction was used in each case to correct for multiple testing of bulk RNA-sequencing datasets (65 tissues) as well as for all single-cell RNA-sequencing datasets (67 cell types), respectively.

### Partitioning heritability of chromatin signatures

We used stratified LD score regression^72,73^ to test for enrichment of endocranial globularity SNP-heritability in annotations of tissue-specific chromatin signatures reflecting regulatory and/or transcriptional activity. The annotations used here were all based on data from the Roadmap Epigenomics project and ENTEX, as described by Finucane et al.^72^. Bonferroni correction was used to correct for multiple testing of 489 annotations.

### Analyses of contributions of Neandertal admixture

We used stratified LD score regression to determine if annotations of the human genome tagging signatures of Neandertal ancestry show enrichment or depletion of the total SNP-heritability. We estimated the contribution of evolutionary annotations associated with Neandertal ancestry, by using recently published and enhanced genome annotations^78^. These annotations were chosen based on their relevance for the timing of the evolution of endocranial globularity in our ancestors (i.e. after the Neandertal-*Homo sapiens* split), and the need for sufficient SNP-coverage to allow robust partitioned heritability analysis. Based on these criteria, the following annotations were used: 1) Neandertal introgressed alleles (and SNPs in perfect LD (*r*^2^ = 1)), short genomic regions of Neandertal DNA which introgressed in to varying degree into the genome of some present-day humans as a result from interbreeding events^74^; and 2) archaic deserts, long stretches of DNA that are significantly depleted of Neandertal sequences in modern human populations^75^. Both annotations were controlled for the baseline LD v2.2 model.

### MAGMA gene-set analysis with custom evolutionary gene-sets

Two curated genome annotations with high evolutionary relevance were unsuitable for heritability partitioning due to low SNP-coverage and were assessed instead via MAGMA gene-set analysis^79^. These annotations were: 1) Ancient Selective Sweeps, haplotypes that reached fixation after the split with archaic humans (∼450-750 kya), but before the differentiation of modern human populations (∼100-120 kya)^76^, 2) Anatomically Modern Human-derived Differentially Methylated Regions (AMH-derived DMRs), regions with unmethylated state in ancient and modern humans, but methylated in Neandertals and Denisovans, reflecting chromatin-state differences that emerged between AMHs and archaic humans after they split ∼450-750 kya^77^. We listed genes that fall within ±1 kb of each loci locus tagged by one of these two annotations, and filtered for protein coding genes using NCBI’s hg19 genome annotation ^135^. We then performed gene annotation by integrating SNP locations from GWAS summary statistics and gene locations from NCBI hg19 genome annotation. We followed this with a gene-based analysis using SNP p-values and 1000 Genomes Phase 3 European Panel^133^, and finally applied the gene-set analysis using results from gene annotation, gene analysis, and the two evolutionary gene-sets that we curated. Bonferroni correction was used to correct the enrichment p-value threshold for two tests.

## Data availability

Imaging data across the adult lifespan are available from CamCAN (http://www.mrc-cbu.cam.ac.uk/datasets/camcan). Neuroimaging and genotype data used for GWAS are available from UK Biobank (https://www.ukbiobank.ac.uk). GWAS summary statistics of endocranial globularity will be made freely available through the GWAS Catalog (https://www.ebi.ac.uk/gwas/).

## Code availability

Coordinate-measurements on endocasts, and scripts used for the analyses are available on the project GitLab repository (https://gitlab.gwdg.de/barmol/globularity).

## Acknowledgments

This research was conducted using the UK Biobank resource under application no. 16066 with CF as the principal applicant. Our study made use of brain imaging derived phenotypes and pre-processed imaging data generated by an image processing pipeline developed and run on behalf of UK Biobank. Data collection and sharing for this project was provided by the Cambridge Centre for Ageing and Neuroscience (CamCAN), with funding provided by the UK Biotechnology and Biological Sciences Research Council (grant number BB/H008217/1), together with support from the UK Medical Research Council and University of Cambridge, UK. SEF is a member of the Center for Academic Research and Training in Anthropogeny (CARTA). We thank all curating institutions for access to CT data of fossil and extant human crania, and the Max Planck Institute for Evolutionary Anthropology Leipzig (Dept. of Human Evolution) for CT scanning. We further thank Giacomo Bignardi for insightful discussions which helped to shape the ethical workup of this study. BM, GA, EE, DS, CF and SEF were supported by the Max Planck Society. EE is also supported by the Dutch Research Council (NWO; VI.Veni.202.072), and PG is also supported by the Evolution of Brain Connectivity Project (M.IF.A.XXXX8103).

## Author contributions

Conceptualization: BM, EE, GA, CF, PG, SEF Resources: CF, PG, SEF

Methodology: BM, EE, GA, DS, PG, SEF Data analysis: BM, GA, PG

Writing – original draft: BM, PG

Writing – review & editing: BM, EE, GA, DS, CF, PG, SEF

## Competing interests

Authors declare that they have no competing interests. Supplementary Information is available for this paper

Correspondence and requests for materials should be addressed to Philipp Gunz or Simon Fisher

