## Supplementary Material for "Imaging genomics reveals genetic architecture of the globular human braincase"

**This file includes:**

Figs. S1 to S6
Tables S1 to S11 (supplied in additional excel table due to table size)
Supplementary references

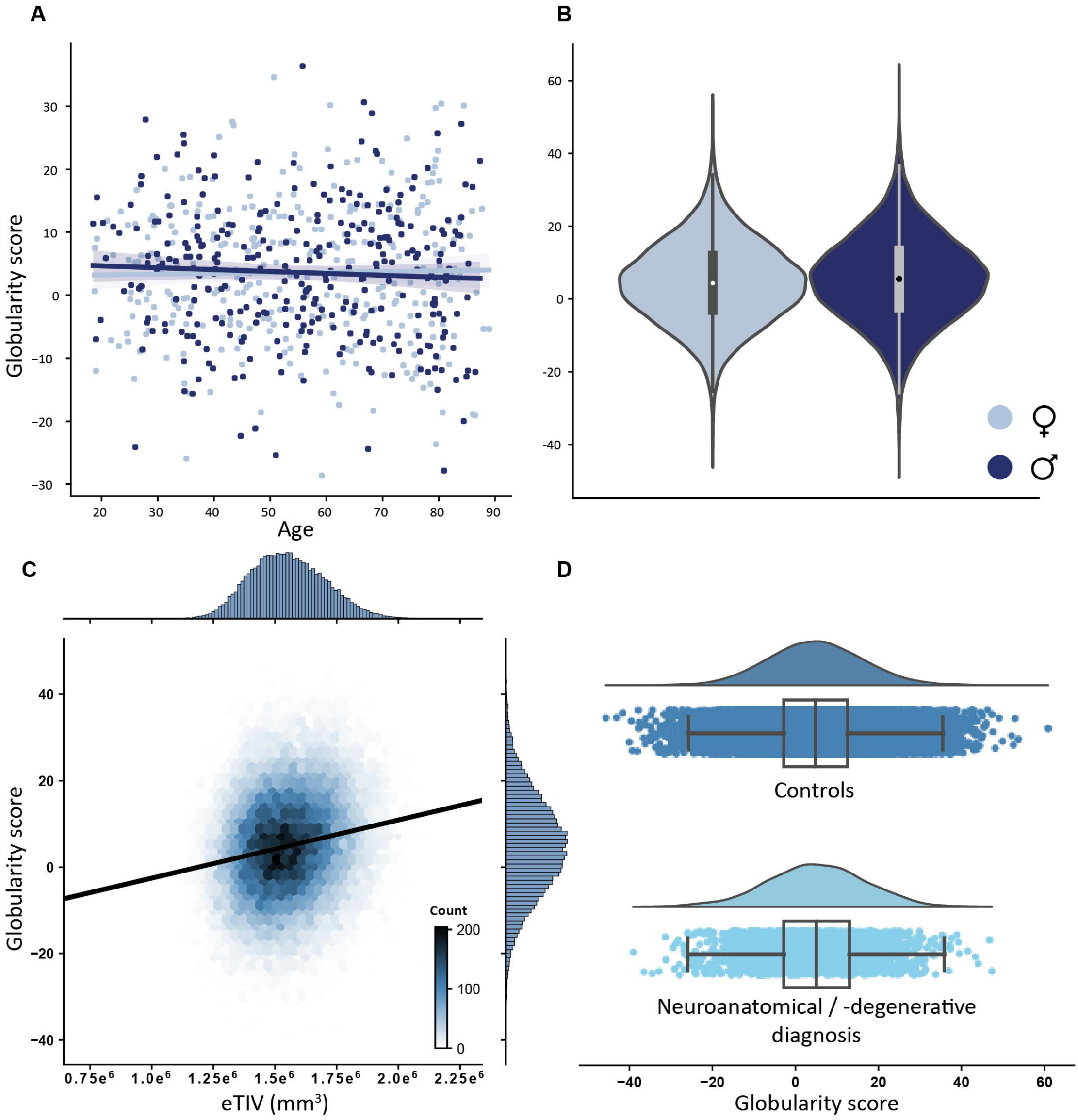


**Fig. S1. Phenotypic assessment of present-day human endocranial globularity. (A)** Relationship between globularity scores and age across the adult lifespan in healthy present-day humans stratified by sex (CamCAN, N = 644; $N_{female}$ = 326, $N_{male}$ = 318); linear regression fit and univariate kernel density estimation curves are added for both males and females separately. **(B)** Violin plot (kernel density estimate) of globularity scores in UK Biobank stratified by sex ($N_{female}$ = 17,803; $N_{male}$ =16,148). **(C)** Relationship between globularity scores and estimated total intracranial volume (eTIV; UKB data field 2652, N = 33,577); hexbin count and linear regression fit are added; histograms depict the marginal distribution of each variable opposite the respective axes with joint distribution highlighted in the middle as (hexbin) scatterplot. **(D)** Distribution of globularity scores for participants diagnosed with a neuro-anatomical / -degenerative disorder (N = 2,338) and control participants (N = 31,613). In the raincloud plots (*1*) the half-violin plots highlight the data distribution, where the adjacent boxplots indicate the 25th and 75th percentiles and whiskers represent 1.5 times the interquartile range. Individual data points are jittered to aid visualization. For A-C, data shown here are unadjusted, but note that the relevant covariates were included in the respective statistical association tests described in the Methods section.

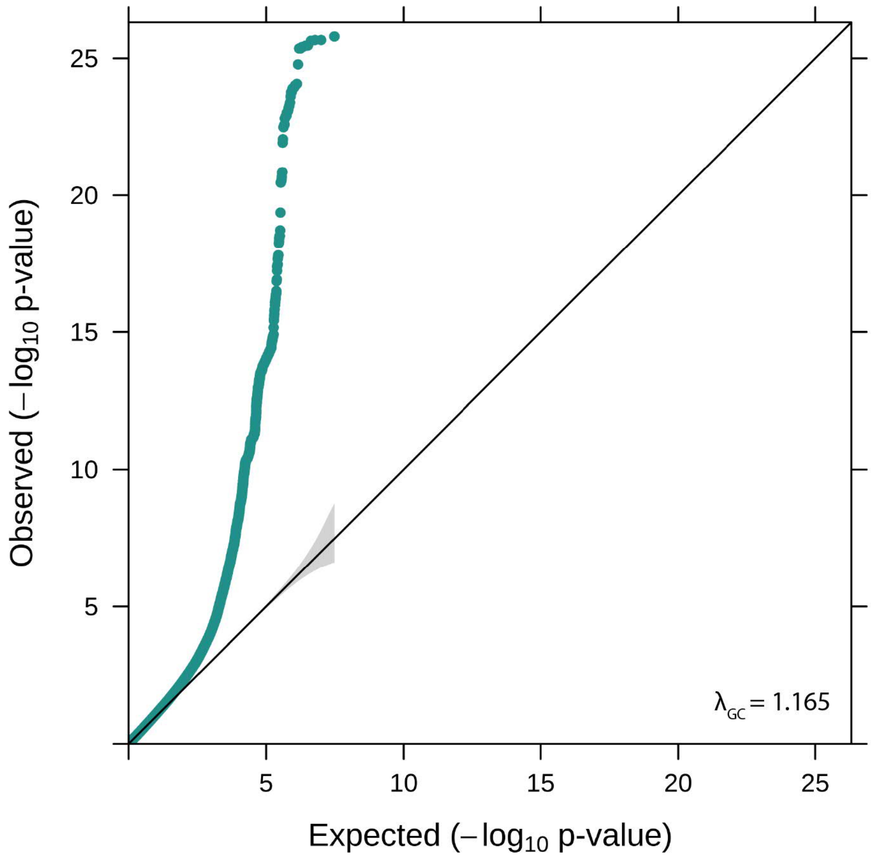


**Fig. S2. Quantile-quantile plot of GWAS for endocranial globularity.** The gray-shaded area represents the 95% confidence intervals under the null hypothesis.

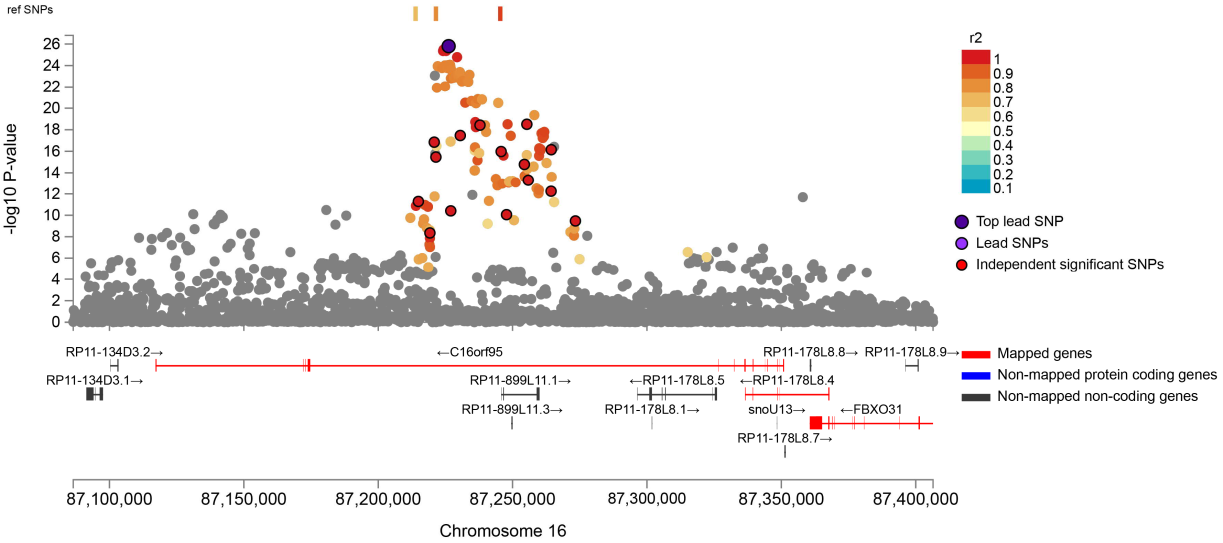


**Fig. S3. Locus zoom plot showing GWAS signals for rs9933149 and SNPs in linkage disequilibrium**.
Locus zoom plot from FUMA, where colours indicate linkage disequilibrium with rs9933149 based on the European 1000 Genomes Project reference data. Overlapping mapped and non-mapped genes are shown below, with prioritized genes highlighted in red.

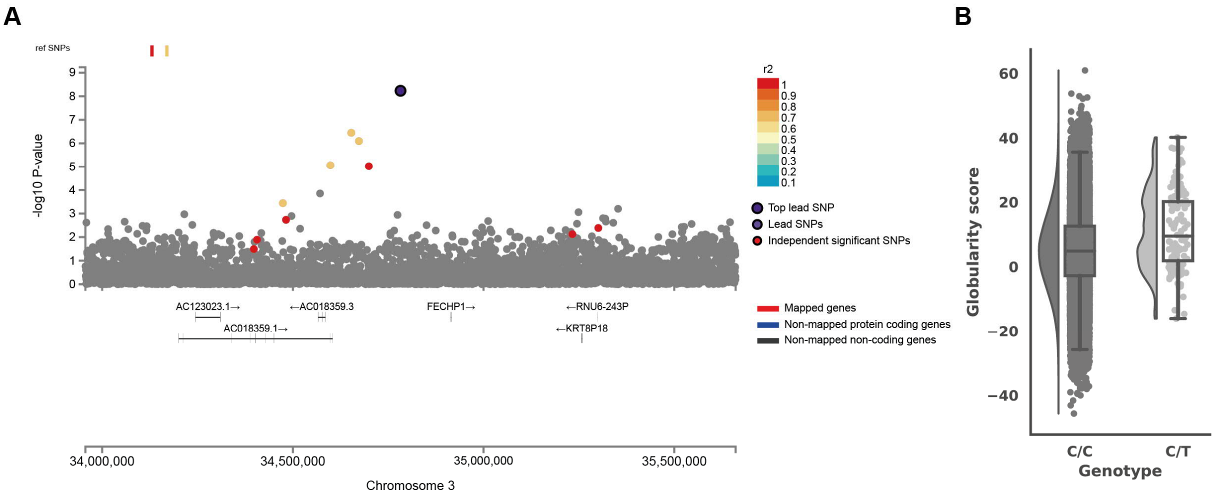


**Fig. S4. High effect size variant rs529711487. (A)** Locus zoom plot of rs529711487 from FUMA, where colours indicate linkage disequilibrium with rs529711487 based on the European 1000 Genomes Project reference data. Overlapping mapped and non-mapped genes are shown below. **(B)** Raincloud plot of uncorrected globularity scores per genotype. All data points are shown; boxes represent 25th and 75th percentiles; whiskers represent 1.5 times the interquartile range.

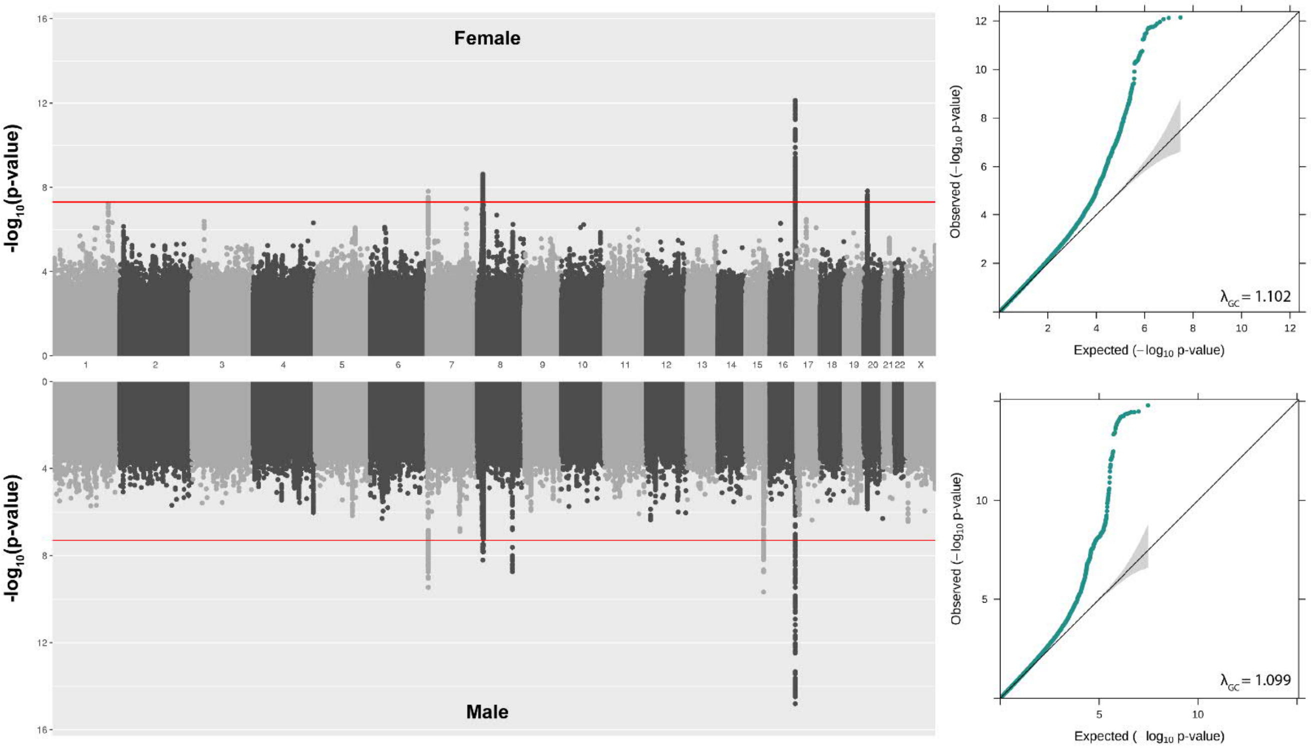


**Fig. S5. Miami and quantile-quantile plots of sex-stratified genome-wide association analyses.** Miami plot of loci associated with interindividual variance in endocranial globularity; the top panel includes only female participants, while the bottom panel includes only male participants. The red horizontal lines denote genome-wide significance. For the quantile-quantile plots, the gray-shaded areas represent the 95% confidence intervals under the null hypothesis.

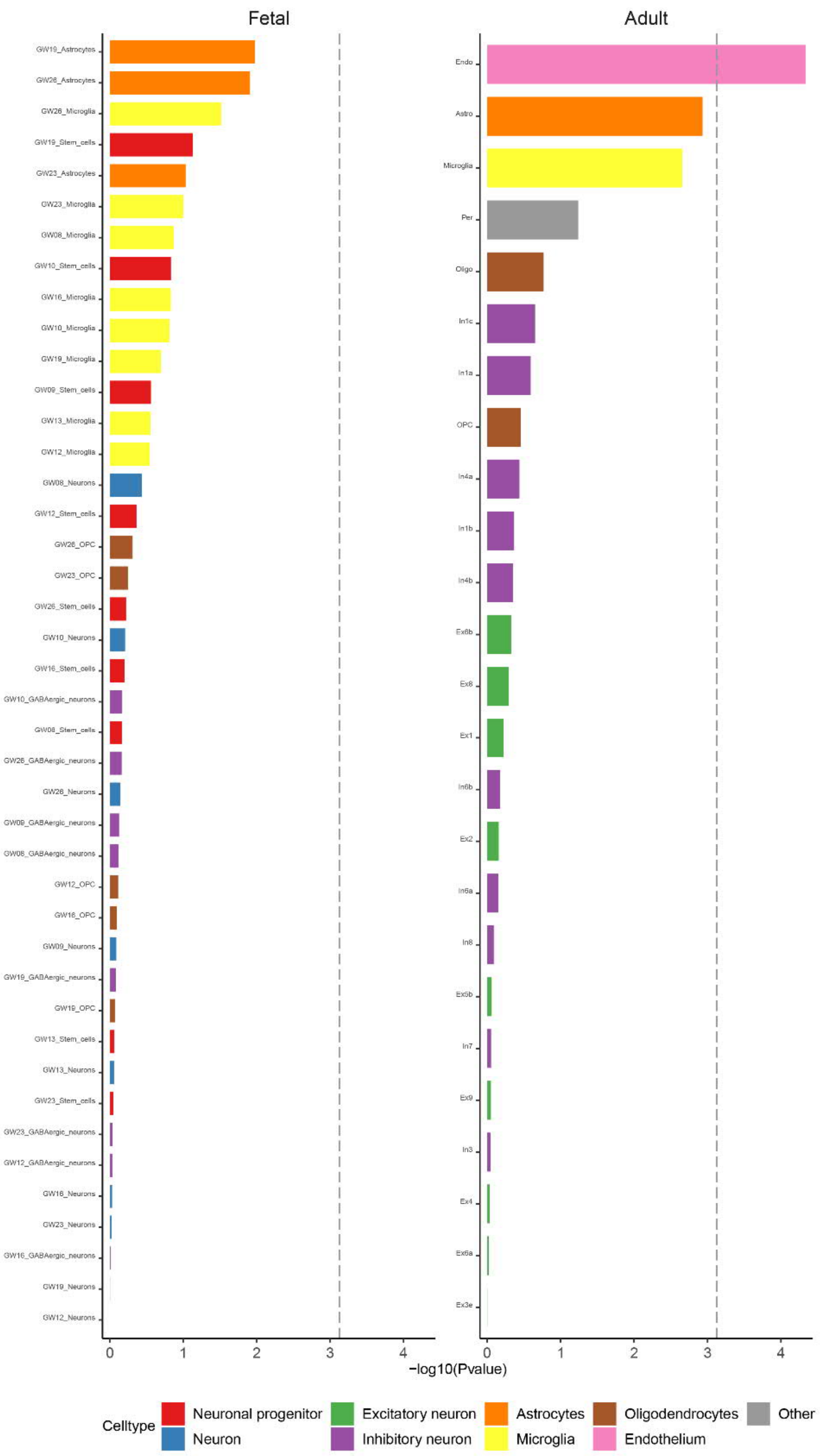


**Fig. S6. Gene-property analysis with MAGMA highlights significant enrichment in astrocytes and endothelial cells in adult brain tissue.** MAGMA gene-property analysis using two single-cell RNA-sequencing datasets of human brain tissue, representing fetal and adult cortex. The graph shows -log-10 p-values on the x-axis and cell types on the y-axis. The dashed line shows the p-value threshold for significant enrichment after Bonferroni correction for testing 67 cell types.

1. M. Allen, D. Poggiali, K. Whitaker, T. R. Marshall, R. A. Kievit, Raincloud plots: a multi-platform tool for robust data visualization. *Wellcome Open Res.* **4**, 63 (2019).
